# Data-Dependent Acquisition with Precursor Coisolation Improves Proteome Coverage and Measurement Throughput for Label-Free Single-Cell Proteomics

**DOI:** 10.1101/2022.10.18.512791

**Authors:** Thy Truong, S. Madisyn Johnston, Kei Webber, Hannah Boekweg, Caleb M Lindgren, Yiran Liang, Alissia Nydeggar, Xiaofeng Xie, Samuel H. Payne, Ryan T. Kelly

## Abstract

The sensitivity of single-cell proteomics (SCP) has increased dramatically in recent years due to advances in experimental design, sample preparation, separations and mass spectrometry instrumentation. Further increasing the sensitivity of SCP methods and instrumentation will enable the study of proteins within single cells that are expressed at copy numbers too small to be measured by current methods. Here we combine efficient nanoPOTS sample preparation and ultra-low-flow liquid chromatography with a newly developed data acquisition and analysis scheme termed wide window acquisition (WWA) to quantify >3,000 proteins from single cells in fast label-free analyses. WWA is based on data-dependent acquisition (DDA) but employs larger precursor isolation windows to intentionally co-isolate and co-fragment additional precursors along with the selected precursor. The resulting chimeric MS2 spectra are then resolved using the CHIMERYS search engine within Proteome Discoverer 3.0. Compared to standard DDA workflows, WWA employing isolation windows of 8-12 Th increases peptide and proteome coverage by ~28% and ~39%, respectively. For a 40-min LC gradient operated at ~15 nL/min, we identified an average of 2,150 proteins per single-cell-sized aliquots of protein digest directly from MS2 spectra, which increased to an average of 3,524 proteins including proteins identified with MS1-level feature matching. Reducing the active gradient to 20 min resulted in a modest 10% decrease in proteome coverage. We also compared the performance of WWA with DIA. DIA underperformed WWA in terms of proteome coverage, especially with faster separations. Average proteome coverage for single HeLa and K562 cells was respectively 1,758 and 1,642 based on MS2 identifications with 1% false discovery rate and 3042 and 2891 with MS1 feature matching. As such, WWA combined with efficient sample preparation and rapid separations extends the depths of the proteome that can be studied at the single-cell level.

## Introduction

The field of single-cell proteomics (SCP) is advancing rapidly due to improvements in experimental design, sample preparation, separations, mass spectrometry (MS) data acquisition and analysis [1–3]. Experimental conditions that are successful for bulk-scale analyses are often suboptimal for single cells and vice versa. For example, during bulk-scale sample preparation, sample cleanup steps are generally necessary to avoid clogging of liquid chromatography (LC) columns and contaminating the mass spectrometer [4, 5], while adsorptive analyte losses during cleanup may be negligible. In contrast, due to much smaller amounts of, e.g., lipids and debris in single cells, a sample cleanup step may not only be unnecessary, but the accompanying sample losses as a percentage of starting material are expected be much more severe [6, 7]. These different requirements have resulted in low-volume, one-pot methods generally being used for low-input samples, while bulk-scale sample preparation continues to employ extensive cleanup.

Similarly, mass spectrometry (MS) acquisition parameters have very different optimum settings for bulk and single cell samples. The large ion flux from bulk samples results in rapid accumulation of a sufficient ion population to produce a productive MS2 spectrum such that tens of MS2 spectra can be collected per second on an Orbitrap instrument [8]. Since the resolution achieved by an orbitrap mass analyzer scales linearly with transient time, a low-resolution orbitrap setting such as 15,000 fwhm at m/z 200 must also be selected to achieve rapid scan rates. However, this low resolution is still sufficient to produce high-confidence peptide-spectrum matches (PSMs) for MS2 spectra that predominantly contain fragment ions from a single peptide [8]. Single-cell samples have a much lower ion flux, and much longer injection times (i.e., up to hundreds of milliseconds) may be required to achieve a sufficient ion population for a productive MS2 scan [9–11]. This results in few collected spectra and low proteome coverage. At these slow scan speeds, a higher resolution (slower) orbitrap resolution setting (e.g., >100,000 at m/z 200) may be used with no impact on cycle time, but this higher resolution is generally excessive and of little benefit for resolving fragments from a single precursor.

The long required injection times and the accompanying high-resolution MS2 spectra that can resolve more complex ion populations point to the intentional precursor coisolation for the identification of multiple peptides as a means of overcoming the slow scan speeds required for most SCP analyses. That is, multiple peptides can simultaneously accumulate for sufficient time in an ion trap, and the fragment ions from these peptides can be readily resolved using a high-resolution orbitrap scan. Indeed, data independent acquisition (DIA)-based SWATH MS [12] steps through large, overlapping precursor isolation windows to generate complex peptide spectra. In contrast, data-dependent acquisition (DDA) software has historically searched for the highest scoring single peptide from a given MS2 spectrum, although approaches have been developed to effectively identify multiple peptides from a single ‘chimeric’ MS2 spectrum [13].

We supposed that the intentional coisolation of multiple precursors using much larger isolation windows (up to 48 Th vs., e.g., 1.6 Th for standard DDA), termed wide-window acquisition (WWA) DDA, would be especially beneficial for SCP performed with Orbitrap instrumentation due to the typical use of long ion accumulation times and their accompanying high-resolution MS2 spectra. To test this, we utilized the recently released CHIMERYS software developed by MSAID (Munich, Germany) and included with Proteome Discoverer 3.0 (Thermo Fisher, Waltham, MA) [14], which can resolve >10 precursors per MS2 spectrum. WWA serves as a hybrid between DDA and DIA (Figure 1), with a specific precursor being isolated for fragmentation (as with DDA), while the wide isolation windows allow for co-fragmentation and analysis of untargeted neighboring precursors (as with DIA). We evaluated WWA for rapid label-free single-cell proteome profiling of single cells prepared using nanoPOTS [6], and found that isolation windows in the range of 8 – 12 Th provide greatest peptide and protein coverage when identification is based solely on MS2 spectra (i.e., without the use of MS1-based feature matching such as the Match Between Runs (MBR) algorithm[15]). These optimized isolation windows agree with the findings of Mayer et al. [16], who have concurrently explored WWA for small aliquots of bulk-prepared samples. Optimized WWA provides a ~30% increase in MS2-identified peptides relative to standard DDA, although peptide and proteome coverage are similar when including MBR identifications. However, due to the increased confidence associated with false discovery rate (FDR)-controlled MS2 identification relative to MBR, it is beneficial to increase the ratio of MS2:MBR-identifications. We also compared WWA to DIA analyzed using MSFragger-DIA with DIA-NN quantification [17], and found WWA provides greater coverage, particularly for faster (20-min) LC separations. Remarkably, in combination with highly sensitive analyses afforded by low-nanoflow LC operated at ~15 nL/min, we have achieved an unprecedented proteome coverage of >3000 protein groups/cell, with nearly 2000 of those proteins identified by tandem mass spectra. The reported workflow thus represents a substantial increase in both sensitivity and acquisition speed for label-free single-cell proteomics.

**Figure 1.**
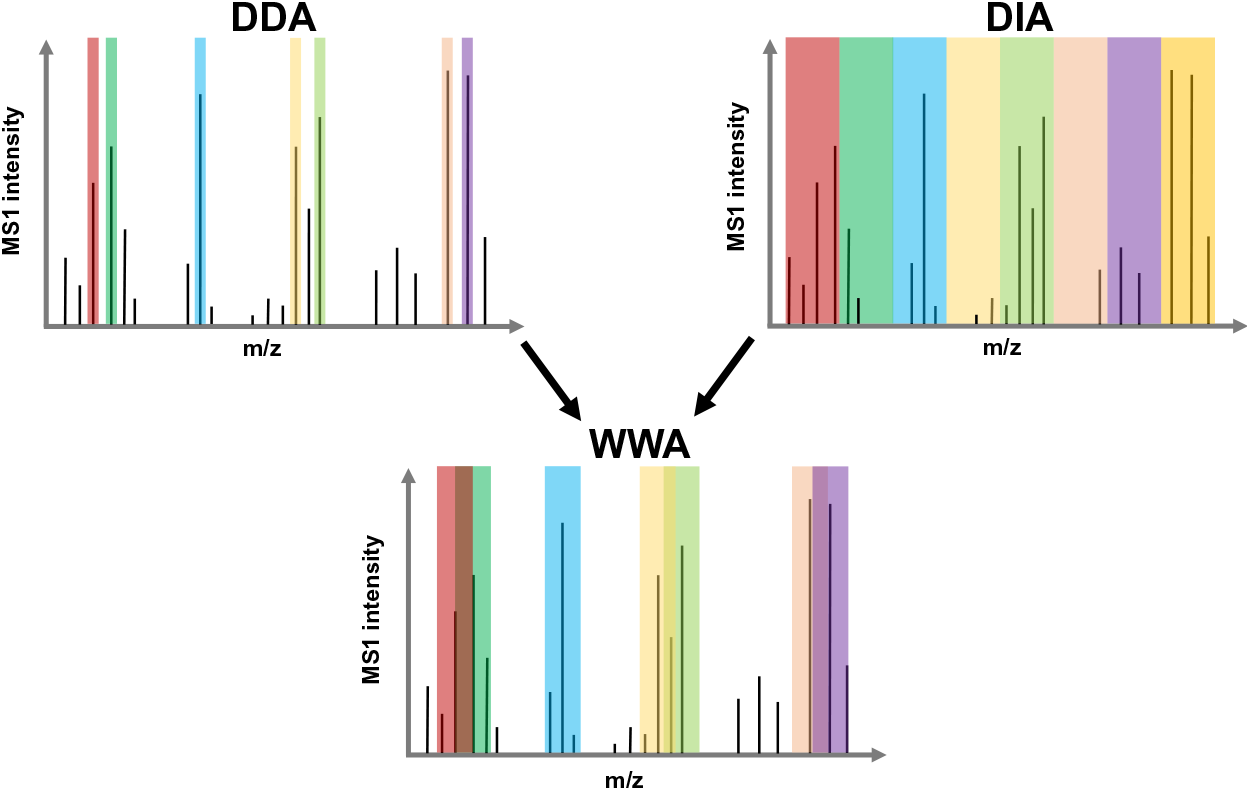
Schematic depiction of precursor isolation using DDA, DIA and WWA.

## Results and Discussion

We anticipated that WWA would increase coverage relative to DDA by quantifying additional peptides that are coisolated with the precursor specifically selected for fragmentation. However, we also anticipated that with a sufficiently large isolation window, the complexity of the resulting spectra may impede identification. As such, we performed screening experiments using single-cell-sized (0.2 ng) aliquots of HeLa digest in which the isolation window was systematically varied from 2 to 48 Th as shown in Figure 2. These analyses were performed across four different maximum injection times/MS2 resolutions ranging from 54 ms/30,000 to 246 ms/120,000. The active chromatographic gradient was ~40 min and the AGC target was set to 1000% such that the maximum injection time rather than AGC target was generally limiting. The number of collected MS2 spectra predictably decreased with increasing maximum injection time/resolution settings, yet the number of PSMs was greatest for the intermediate MS2 resolutions of 45k and 60k (Figure 2A). For larger isolation windows, the number of identified PSMs exceeded and in some cases more than doubled the number of collected MS2 spectra, indicating effective identification of multiple precursors per MS2 spectrum by the CHIMERYS search engine.

**Figure 2.**
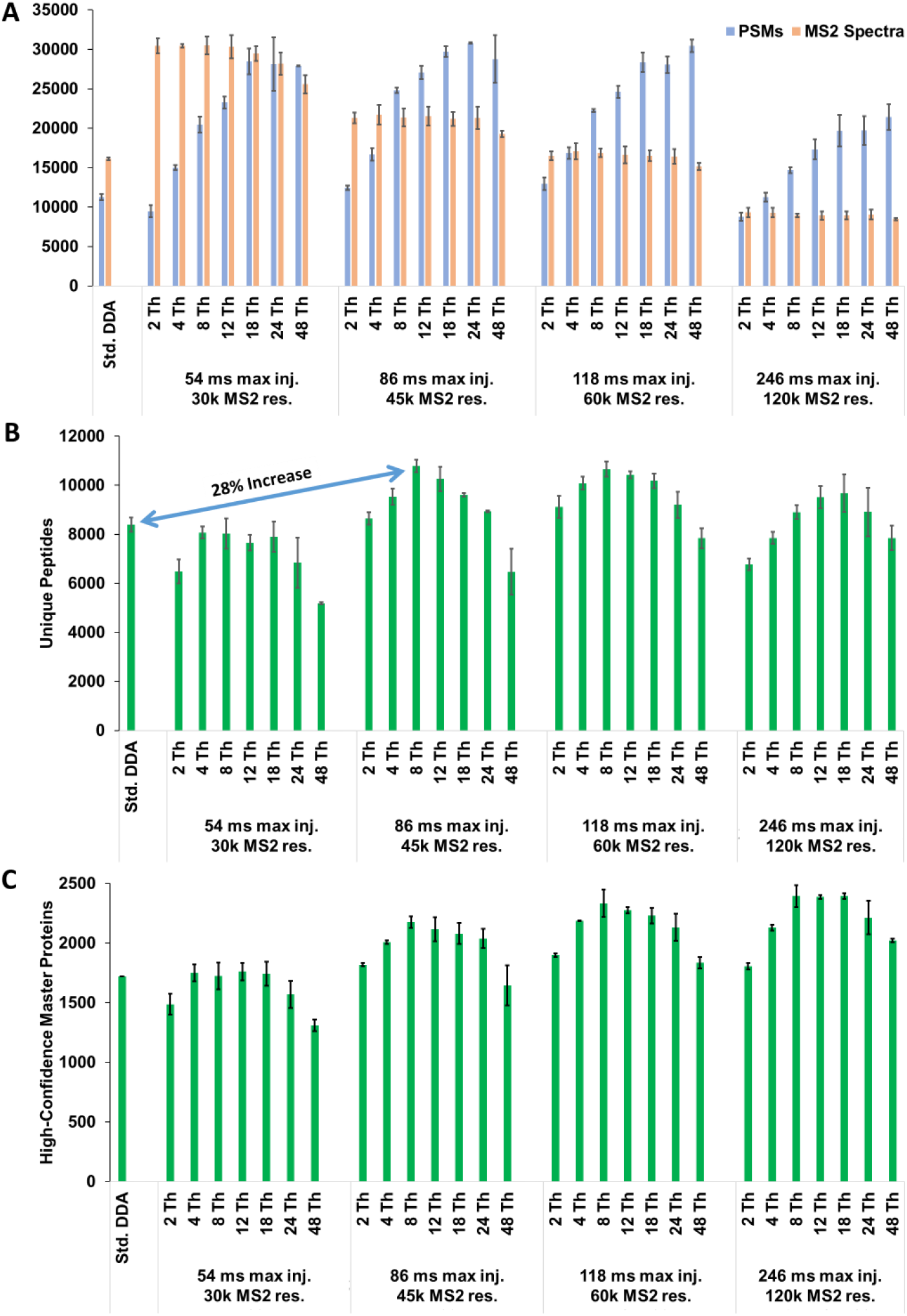
Parameter optimization experiments for 40-min gradients using 0.2 ng aliquots of HeLa digest. A) Number of collected MS2 spectra and PSMs as a function of maximum injection time/resolution and MS2 isolation window. (B) Number of unique peptides identified as a function of MS acquisition settings. (C) Number of high-confidence master proteins (1% FDR). All identifications are based solely on MS2 identification (i.e., no MBR). Std. DDA conditions are listed in Methods. Error bars indicate ± 1 std. dev., and n=2 for all conditions.

However, while >30,000 PSMs could remarkably be identified from single-cell-sized samples, far fewer unique peptides were identified (Figure 2B), indicating that many of these PSMs were redundantly sequenced due to the large and overlapping isolation windows. Still, identifying up to an average of 10,789 unique peptides with 1% FDR and without MBR is ~28% greater than our DDA-based coverage (average 8,399) when using the same separation and MS instrumentation (Figure 2B). Peptide coverage was consistently greatest for isolation windows of 8 or 12 Th, which is in agreement with the findings of Mayer et al. [16], and which points to a compromise between maximizing the number of isolated precursors and avoiding overly complex MS2 spectra. At the protein level, the number of MS2-identified high-confidence master proteins from 0.2 ng aliquots of HeLa digest increased 39% to 2,396 for WWA compared to 1,721 identified on average using standard DDA (Figure 2C).

We also employed MBR, known as Feature Mapper in PD 3.0, using 10 ng aliquots of the same HeLa digest sample as a matching library. We found that MBR substantially increased peptide coverage and that the coverage was essentially independent of MS2 acquisition parameters, as shown in Figure S1. Indeed, 19,478 unique peptides were identified on average across all acquisition conditions including standard DDA, with just 3.9% CV. Similarly, proteome coverage including MBR was highly uniform across all screening conditions: 3,484 high-confidence master proteins were identified on average, with 2.0% CV. Only 171 proteins were identified on average from blank runs, including those identified by MBR, indicating a low degree of column carryover. Given the uniform coverage observed across screening conditions after applying the MBR algorithm, it should be determined whether there are benefits that remain for alternative acquisition strategies such as WWA. To address this, we estimated the false matching rate by generating a mixed-species library comprising human and yeast tryptic peptides in a similar manner to Woo et al. [18]. Among 9,252 total peptides identified by MS2, 40 (0.43%) precursors were matched to nonhomologous yeast peptides. For MBR-based identifications, 483 of 15,070 unique peptides (3.2%) were incorrectly matched to yeast. As such, while transferred identifications from the MBR algorithm greatly increase the depth of attainable proteome coverage, it is beneficial to employ WWA in conjunction with MBR, as it maximizes the ratio of MS2: transferred identifications to reduce the overall FDR.

Shorter LC gradients having reduced peak capacities increase the number of coeluting peptides, which poses a challenge for standard DDA. Since the CHIMERYS search engine can decipher complex, highly chimeric fragmentation spectra, it should also benefit more rapid analyses. We also performed screening experiments for faster separations having ~20-min elution windows (Figure S2) and found 12 Th isolation windows with and 60k MS2 resolution were highly effective in maximizing MS2-identified peptides. We compared proteome coverage between 40-minute and 20-minute gradients using the same acquisition parameters, as shown in Figure 3A. Impressively, MS2-based protein identifications only decreased by 9% for the faster analyses (1953 vs. 2150). With MBR, a 10% reduction in coverage was observed (3160 vs 3524). This indicates that the WWA can accommodate much faster separations with only modest impact on proteome coverage.

**Figure 3.**
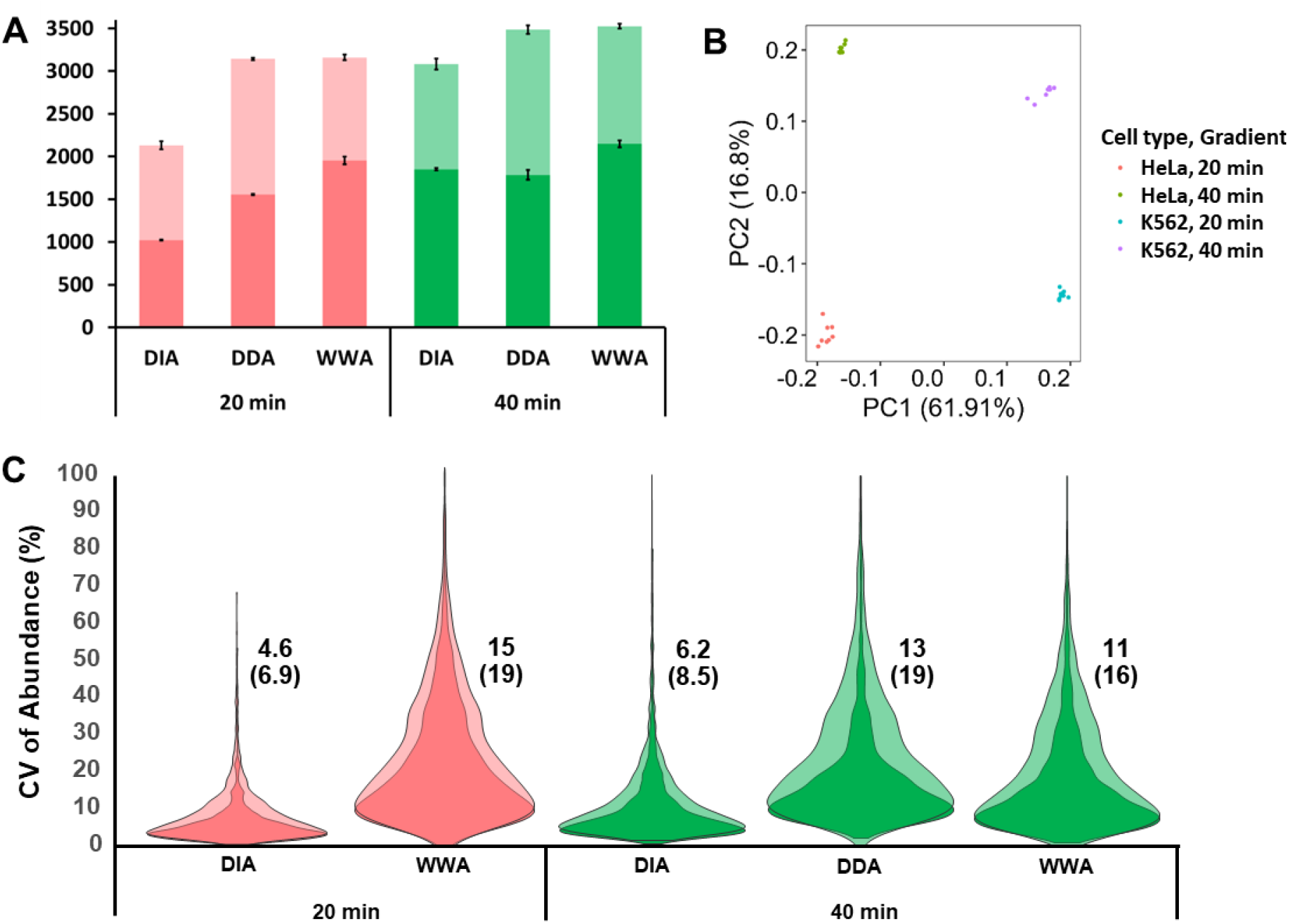
Analysis of 0.2 ng aliquots of protein digest using WWA, Std. DDA and DIA with 20 and 40-min gradients. (A) Proteome coverage using the three data acquisition modes, with MS2-based identification shown in darker shading and additional MBR-identified proteins shown in lighter shading. (B) PCA plot resolving two cell types (HeLa and K562) in PC1, and cells of a given type resolved in PC2 according to gradient length. (C) Violin plots indicating CV of protein abundance for different acquisition modes and gradient lengths, using the same color and shading scheme as (A). Median %CVs for each group are shown next to each plot, with and without MBR (WWA and DDA) or a spectral library search (DIA) shown without and with parentheses, respectively. Eight replicates were obtained in all cases except for Std. DDA in employing 20-min gradients, which had only two replicates.

Given that WWA may be considered a hybrid between traditional DDA and DIA, we sought to also compare its performance to DIA. Using MSFragger-DIA with DIA-NN quantification [17]. We first evaluated various DIA acquisition settings and found no significant difference in coverage (Figure S3). As such, we selected a fixed 50 Th isolation window scanned between 400-800 Th as described by Gebreyesus et al. for further experiments [19]. Results were obtained both with and without employing a spectral library obtained from analyzing 10 ng of HeLa digest. Under all conditions, WWA provided greater proteome coverage (Fig. 3A). The reduction in coverage for DIA when employing faster gradients was much greater than for WWA. As also shown in Figure 3A, proteome coverage for DIA decreased by 45% with library-free search (1021 vs. 1850), and by 31% when a spectral library was employed (2131 vs. 3083).

The ability to differentiate distinct cell types, treatment groups, etc. is of course critical for SCP. We used principal component analysis (PCA) to determine the ability of WWA to cluster technical replicates of HeLa and K562 digest according to cell type for both 20 and 40-min gradients. As shown Figure 3B, HeLa and K562 cell types readily cluster according to cell type along the PC1 axis, which accounts for 62% of variance. Replicates for a given cell type are also well differentiated along the PC2 axis according to gradient length, presumably due to increased coverage for the longer gradients, which accounts for another 17% of explained variance.

We investigated CVs in protein intensities across 8 technical replicates for the different acquisition/analysis methods and gradient lengths. Note that standard DDA experiments employing 20-min LC gradients was excluded from this comparison as only two technical replicates were obtained. The darker shades in Figure 2B indicate no MBR/spectral library, while the lighter shades showing slightly increased median CVs include MBR (DDA and WWA) or spectral library matching (DIA). Median %CVs are shown with and without matching, with matching %CV values provided in parentheses. WWA provided improved the reproducibility of intensity measurements relative to DDA, while DIA provided lower CVs than the other methods. We initially supposed that this was due to increased sampling rates across the narrow chromatographic peaks when using DIA. However, DIA-NN reports an average of 4.07 data points across the FWHM peak for 40-min separations, which is very close to the 4.09 points obtained for the corresponding WWA analyses as obtained by dividing the average FWHM peak width of 6.14 s by the 1.5 s cycle time. Similarly, DIA-NN reports an average of 3.09 points across the FWHM peak for 20 min DIA analyses, which is very similar to the 3.19 points per FWHM peak for WWA, when the average FWHM peak width decreases to 4.78 s. As such, it appears that other differences in assigning intensity values between DIA-NN and PD 3.0 account for the decreased CVs observed with DIA. We also investigated internal consistency in assigning intensity values between WWA and DIA using their respective data analysis algorithms. Figure 4A compares mean intensities observed across 8 replicates between 40-min and 20-min WWA analyses. The assigned intensities are in close agreement, with a correlation coefficient of 0.91 (Fig 4A). In contrast, DIA provided a more modest correlation of 0.74 between slow and fast analyses (Fig. 4B). Comparing intensities between 40-min WWA and DIA analyses resulted in a correlation coefficient of 0.78 (Figure 4C), with differences arising presumably from MS1-based identification of WWA vs. use of MS2 intensities for DIA.

**Figure 4.**
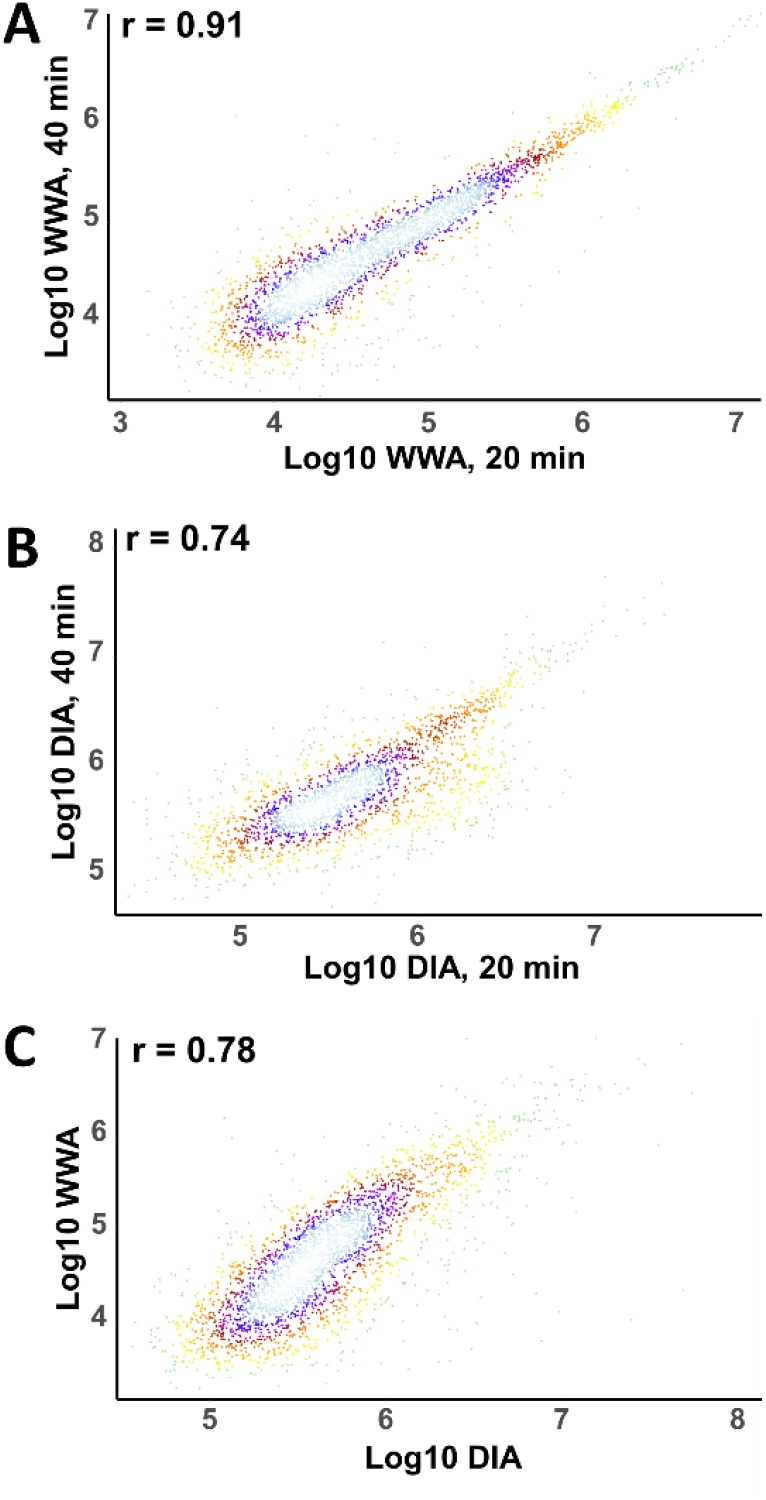
Comparison of protein abundance values obtained using different data acquisition and analysis schemes. Additional description is in the text.

Having established the achievable proteome coverage and quantitative reproducibility of WWA with technical replicates of single-cell-sized aliquots of protein digest, nanoPOTS-prepared single HeLa and K562 cells were then analyzed. For single cells, we maintained the same maximum injection and MS2 resolution settings of 118 ms and 60,000. However, we narrowed the isolation window from 12 to 8 Th, anticipating that increased contaminant levels in single cells may limit ion accumulation times. Ten HeLa cells and 7–10 K562 cells were analyzed using each of the acquisition modes (DIA, DDA and WWA) with 40 min LC gradients. As with the aliquots, WWA provided the greatest proteome coverage, both with and without MBR/spectral library (Figure 5A). Without MBR or a spectral library, an average of 1,758 proteins were identified from HeLa cells by WWA, compared to 1,350 by DIA and 1,312 by DDA, providing a respective increase of 30% and 34%. Including transferred identifications from MBR or a spectral library, WWA proteome coverage was 24% greater than DIA (3,042 vs. 2,463), yet only 3% greater than DDA (3042 vs. 2943). These findings were thus consistent with those resulting from the analysis of bulk-prepared digests. As shown in Figure 5B greater median CVs were observed for single cell experiments relative to aliquots of bulk-prepared digest, as expected due to biological variability, and DIA continued to show less variability in observed protein intensities than the other acquisition modes. Similar proteome coverage was observed for HeLa and K562 cells, and these two cell types were readily differentiated by PCA along PC1 (Fig. 5C).

**Figure 5.**
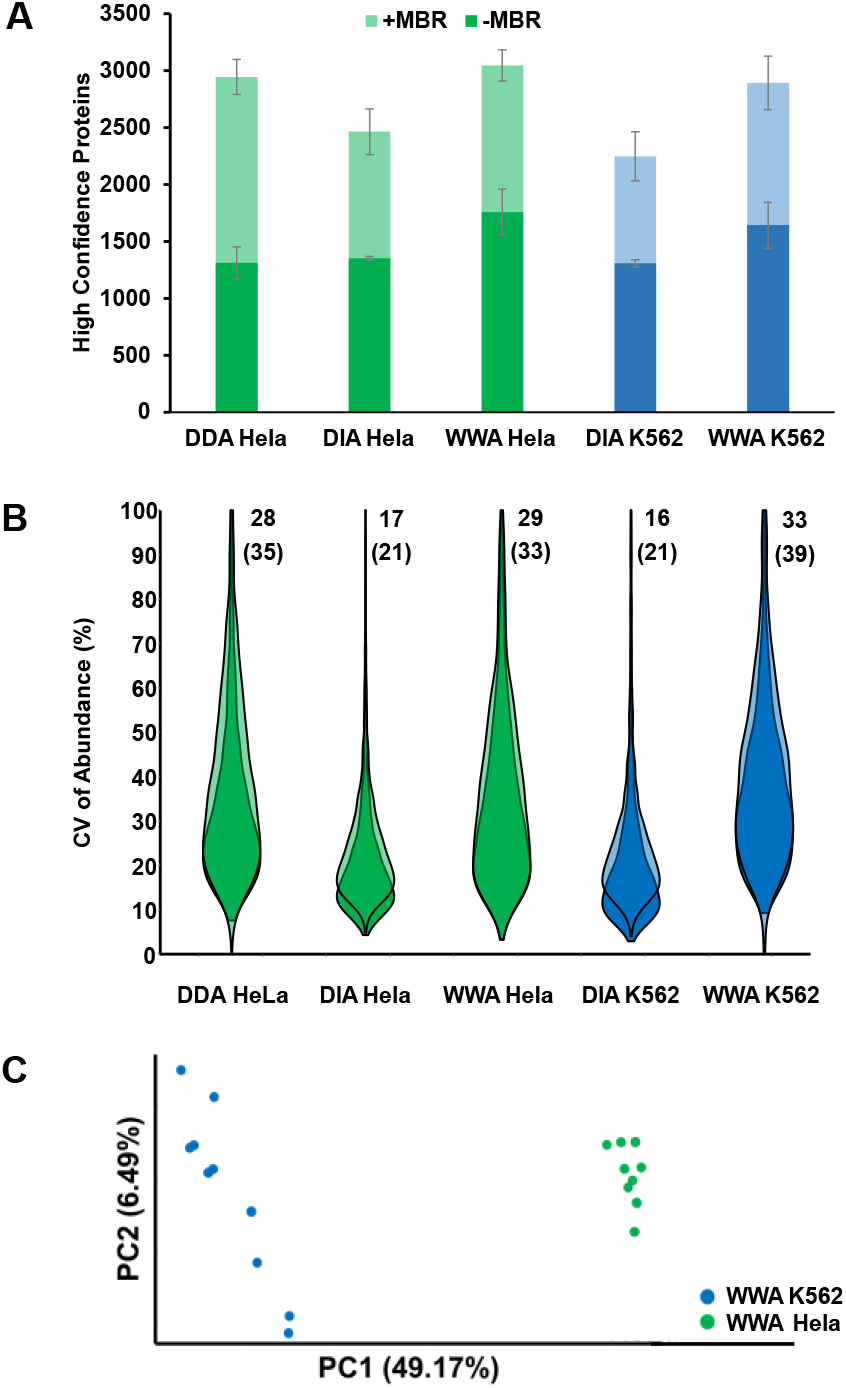
Analysis of nanoPOTS-prepared single HeLa and K562 cells. (A) Proteome coverage using the three data acquisition modes, with MS2-based identification shown in darker shading and additional MBR-identified proteins shown in lighter shading. (B) Violin plots indicating CV of protein abundance for different acquisition modes and gradient lengths, using the same color and shading scheme as (A). Median CVs for each group are shown next to each plot, with and without MBR (WWA and DDA) or a spectral library search (DIA) shown without and with parentheses, respectively. (C) PCA plot resolving HeLa and K562 cells. n = 10 for each cell type/analysis method.

To contextualize the increased depth of discovery afforded by the present method, we compared the quantifiable proteins from this study to a previous study by Bekker-Jensen et al. [20], which used bulk samples and extensive fractionation to calculate the protein copies per cell for >12,000 proteins in HeLa cells. We plotted the depth of discovery for all 3,840 detectable proteins observed in our analysis of single HeLa cells according to their copy number per cell as determined previously [20] (Figure 6A). Our single-cell data identifies more than half of the proteins at ≥10^5^ copies per cell, while protein coverage substantially drops off by 10,000 copies per cell. The median abundance of identified proteins is 256,626 or 426 *z*mol. copies per cell. Several low abundance proteins are well represented in our dataset, including 111 present at 6000 copies per cell (~10 zmol) or fewer, although their presence in the single cells may be at higher levels than were observed in the bulk study [20]. 25% of quantified proteins are present at 88,945 copies per cell (148 zmol) or less. In comparison to our lab’s prior efforts [11], we observe approximately one order of magnitude increase in depth. A large portion of proteins in HeLa cells were calculated by Bekker-Jensen to be present at 1,000 to 10,000 copies per cell. Detection of these ultra-low abundance proteins with LC-MS methods is very challenging, yet with additional advances in MS sensitivity and improvements in ionization efficiency afforded by low-flow separations such as open-tubular LC [21], these species may also become amenable to profiling by single-cell mass spectrometry.

**Figure 6.**
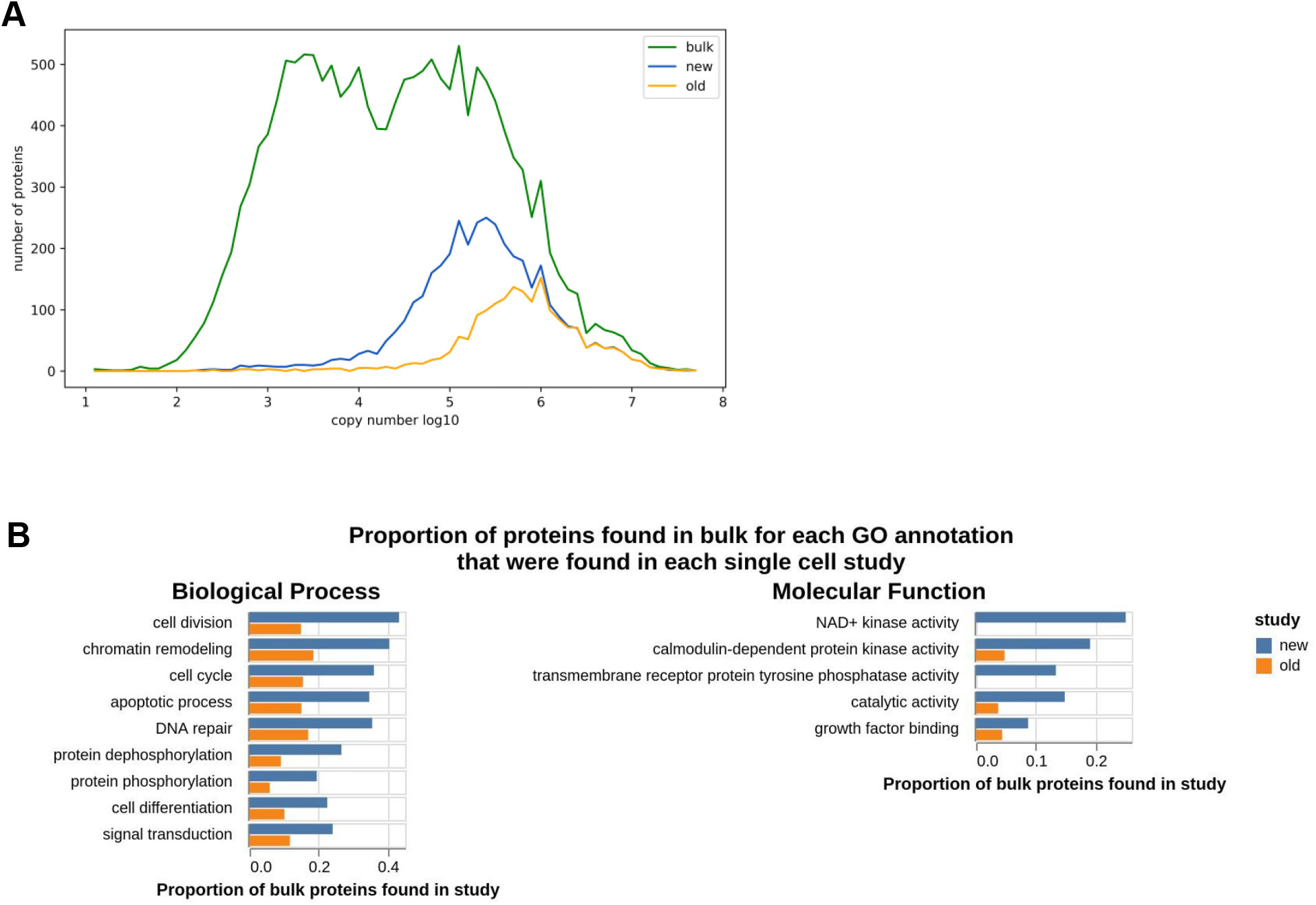
Depth of coverage in wide-window acquisition compared to a traditional LFQ single-cell dataset. (A) The number of proteins detected based on the copy number for three datasets: a bulk proteome study [20], the new wide-window acquisition data (new), and data from our previous LFQ single-cell (old) [11]. (B) A Gene Ontology analysis shows that the wide-window acquisition is able to achieve greater depth for proteins whose functions are important to cancer development and cellular physiology.

With more than twice the number of quantified proteins from our previous analytical methods, we sought to understand the new biological processes that could be observed in our data. Our goal was to investigate whether these newly quantifiable proteins represent a meaningful addition to our ability to describe and characterize cellular physiology, as opposed to uncharacterized proteins or those with ambiguous function. Therefore, we classified proteins according to their Gene Ontology Biological Processes (Table S1). Select biological processes are shown in Figure 6B. For example, several important functions relating to cancer physiology show a significant increase in proteins now detected such as cell division, chromatin remodeling, apoptosis, DNA repair, and cell cycle (i.e., cell cycle regulation). Moreover, proteins involved in the essentials of cellular physiology such as protein phosphorylation and de-phosphorylation, signal transduction and cellular differentiation have also substantially improved in coverage.

## Conclusions

Here we report an improved workflow for label-free single proteomics that incorporates each of the unique advantages of the nanoPOTS sample preparation platform, ultra-low-flow liquid chromatography, and a newly developed WWA data acquisition WWA. Compared to our previous DDA, the new WWA acquisition resulted in increase coverage of up to ~2150 proteins (~3524 with MBR) from 0.2 ng aliquots Hela digest, respectively. WWA also increases the overall throughput by accommodating much faster gradients with only a modest reduction (9%) in coverage. Compared to different DIA variants, WWA coverage outperformed under all conditions, especially at faster gradients. Utilizing all the advantages of the new workflow, we achieved an average of 1758 and 1642 proteins for single Hela and K562 cells without matching (3042 and 2891 with matching). Further investigation using published bulk samples as a reference, we also compared the new single-cell WWA workflow to the previously reported DDA result. With the addition of 3464 annotations from the bulk study found only in the new single-cell study, new cellular functions and biological processes were also observed, as well as an increase in those already detected.

## Methods

### Sample Preparation

Pierce™ HeLa protein digest standard and formic acid were purchased from Thermo Fisher Scientific (Waltham, MA). K562 and yeast digest standards were from Promega (Madison, WI). Mobile phase A (0.1% formic acid in water) and mobile phase B (0.1% formic acid in acetonitrile) were respectively prepared from LC-MS grade water and acetonitrile purchased from Honeywell (Charlotte, NC). The HeLa and K562 digest standards were reconstituted to a final concentration of 200 ng/μL in 100 μL of mobile phase A to form stock solutions. For experiments, the stock solutions were further diluted in mobile phase A to 10 ng/μL and 0.2 ng/μL. To create a mixed-species proteome, yeast and HeLa digests were combined at 8:2 HeLa:yeast at a total concentration of 10 ng/μL. HeLa and K562 cells (ATCC, Manassas, VA) were cultured and harvested as described previously [22]. Following pelleting and removal of cell media, cells were resuspended in phosphate-buffered saline (PBS) to reach a concentration of ~300,000 cells/mL. The cellenONE X1 (Cellenion, Lyon, France) was used for single-cell isolation and reagent dispensing for nanoPOTS sample processing [22] and dried on-chip prior to analysis.

### Separations

Analytical and micro-solid-phase-extraction (SPE or trap) columns were prepared in-house [10] using Dr. Maisch (Ammerbuch, Germany) ReproSil-Pur C18 media having 1.9 μm diameter and 120 Å pore size. Columns were packed in 20-μm-i.d. × 30-cm-long fused silica capillary (Polymicro, Phoenix, AZ). Trap columns were 50-μm-i.d. × 5-cm. Both columns were fritted using Kasil Frit Kit (Next Advance, Troy, NY). Trap columns were fritted on both ends to enable bidirectional flow. Capillary ends were polished using the Capillary Polishing Station (ESI Source Solutions, Woburn, MA). A 10-μm-i.d. chemically etched nanoelectrospray emitter from MicrOmics Technologies (Spanish Fork, UT) were connected to the analytical column via a PicoClear union (New Objective, Woburn, MA).

Aliquots of bulk-prepared protein digest were analyzed from glass vials using an Ultimate 3000 nanoLC system (Thermo Fisher) modified with a 10-port valve as described previously [23]. Sample was loaded onto the SPE at ~ 0.3 μL/min for 10 min at 1% mobile phase B before the valve was switched to deliver the sample onto the analytical column. The flowrate through the analytical column was ~15 nL/min. Electrospray potential (2.2 kV) was applied to the Nanovolume union (VICI, Houston, TX) upstream of the analytical column. Single cell samples were analyzed directly from nanowell chips via a custom autosampler as described previously [24]. For 40-min active gradients, mobile phase B was increased from 1 to 2% in 1 min, 2 to 8% in 5 min, 8 to 15% in 15 min, 15 to 20% in in 9 min and 20 to 25% in 6 min, and 25 to 45% in 10 min. For column washing and regeneration, mobile phase B was increased from 45 to 80% B in 5 min, stepped to 90% for 5 min, stepped to 1% and held for 25 min. For 20-min active elution gradients, the following steps were modified: mobile phase B was ramped from 8 to 15% in 7.5 min, then to 20% in 4.5 min and to 25% in 3 min.

### MS Acquisition

The LC column/emitter assembly was interfaced with an Orbitrap Exploris 480 mass spectrometer (Thermo Fisher) via the Nanospray Flex Ion Source. The temperature of the ion transfer tube was set to 200 °C and the RF lens setting was 50%. For MS1, the Orbitrap resolution was set to 120,000 (m/z 200) with the normalized AGC target set to 300%. The scan range was between 375-1575 m/z and the maximum injection time was set to 118 ms. To trigger MS2 for all experiments, the precursor intensity threshold was set to 5.0E3, charge state was 2 to 4, dynamic exclusion was 20s for 20 min LC gradients and 25s 40 min gradients. The cycle time was 1.5 s. For standard DDA, the isolation window was 1.6 Th, the HCD collision energy was 30%, and the the MS2resolution was respectively set to 15,000 and 60,000 for 10 and 0.2ng HeLa/K562 protein digest standard. The maximum injection time for 10 and 0.2 ng was 22 and 118ms, respectively, and the AGC target was 200%. For WWA, a combination of settings for the isolation windows, resolution, and injection time were evaluated. The AGC target was set to the maximum 1000% for all experiments. The isolation widths were 2, 4, 8, 12, 18, 24 and 48 Th. The MS2 resolution was 30,000, 45,000, 60,000 and 120,000 with corresponding 54 ms, 86 ms,118 ms and 246 ms injection times. For single cell samples, we selected an isolation window of 8 Th for WWA with 60,000 MS2 resolution and 118 ms injection time.

For DIA experiments, the precursor scanning range was from 400 to 800 m/z. A fixed window of 50 m/z increment as well as SWATH acquisitions are detailed in Supplemental Table 1. Window overlap was 1. HCD collision energy was 30%. Resolution of 60000, injection time of 118ms and AGC target of 1000% for 0.2 ng standard and single cell experiments. For 10 ng DIA library, 30000 resolution, 54ms injection time and AGC target of 1000% were set.

### Data Analysis

For DDA and WWA experiments, raw files were searched using Proteome Discoverer (PD) (Thermo Scientific, version 3.0.1.13) with the CHIMERYS identification node using prediction model inferys_2.1_fragmentation as default settings. Database search included human (Uniprot version 2022-8-11) and yeast (uniport, version 2022-8-11) as well as common contaminants (PD, version 2018-10-26). Enzyme was set as Trypsin with maximum 2 missed cleavages. Other parameters included peptide lengths of 7-30 amino acids, a maximum of 3 modications, and charges between 2 and 4. Fragment mass tolerance was 20 ppm. Carbamidomethyl (C) was fixed as a static modification by the CHIMERYS software. Results were filtered with Percolator node at 1% FDR. For MBR, retention time tolerance was set at 0.25 min with a mass tolerance of 5 ppm.

For DIA experiments, FrapPipe (version 1.8) and DIA-NN (version 1.8.1) were used with default DIA_Speclib_Quant workflow. The spectral libraries were generated by FragPipe with RT calibration set to ciRT. Quantification was performed with DIA-NN with Robust LC (high precision) as quantification strategy and MBR enabled. Database search was also human (version 2022-8-11) with added decoys and common contaminants. FDR was set to 1%.

For CV calculations, Pearson r calculations, and quantifiable proteins, all protein data were imported into the R programming language from the “…_Proteins.txt” files exported from PD or the “…_pg_matrix.tsv” exported from DIA-NN. Medium and low confidence proteins were removed, as were contaminants, and for PD any proteins with less than 2 unique peptides. Next, these proteins were sorted according to gradient lengths and sample size, normalized to the median in the separate groups, and filtered for missing values < 33%. For MBR missing values, this meant any without “High” or “Peak Found” proteins in PD files, and just those without “High” for MS2 quantifiable proteins. DIA-NN used only valid values in either the data analyzed with or without a 10-ng spectral library. Pearson correlations used log-10 transformed mean abundances of quantifiable proteins. For PCA, k nearest neighbors was used to impute missing values with a k = 5, and the prcomp function was used in R. All figures were exported to Microsoft PowerPoint for further formatting.

## Data Availability

The mass spectrometry proteomics data have been deposited to the ProteomeXchange Consortium via the PRIDE [25] partner repository with the dataset identifier PXD037527.

## Acknowledgements

The research reported in this publication was supported by the National Cancer Institute and the National Institute of General Medical Sciences of the National Institutes of Health under award numbers R21CA272326 and R01GM138931. The content is solely the responsibility of the authors and does not necessarily represent the official views of the National Institutes of Health.

## Notes

### Competing Interest Statement

RTK has a financial interest in MicrOmics Technologies, which provided nanoelectrospray emitters for this study.

