## Supplementary Information for "Data-Dependent Acquisition with Precursor Coisolation Improves Proteome Coverage and Measurement Throughput for Label-Free Single-Cell Proteomics"

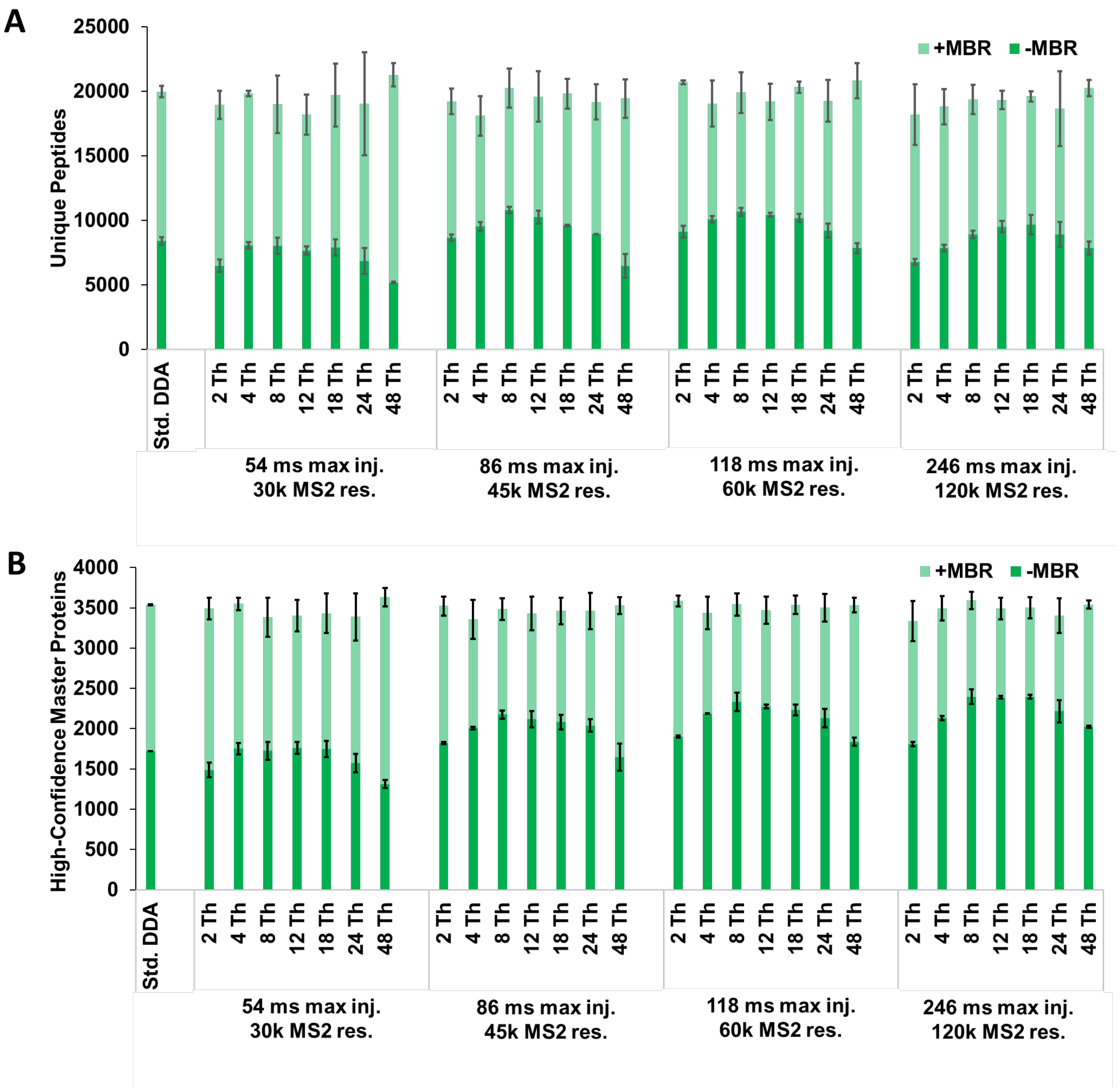


**Figure S1. Parameter optimization experiments for 40-min gradients using 0.2 ng aliquots of HeLa digest with MBR.** (A) Number of unique peptides identified as a function of MS acquisition settings. (B) Number of high-confidence master proteins (1% FDR). All identifications are based on MS2 identification and MBR. Std. DDA conditions are listed in Methods. Error bars indicate ± 1 std. dev., and n=2 for all conditions.

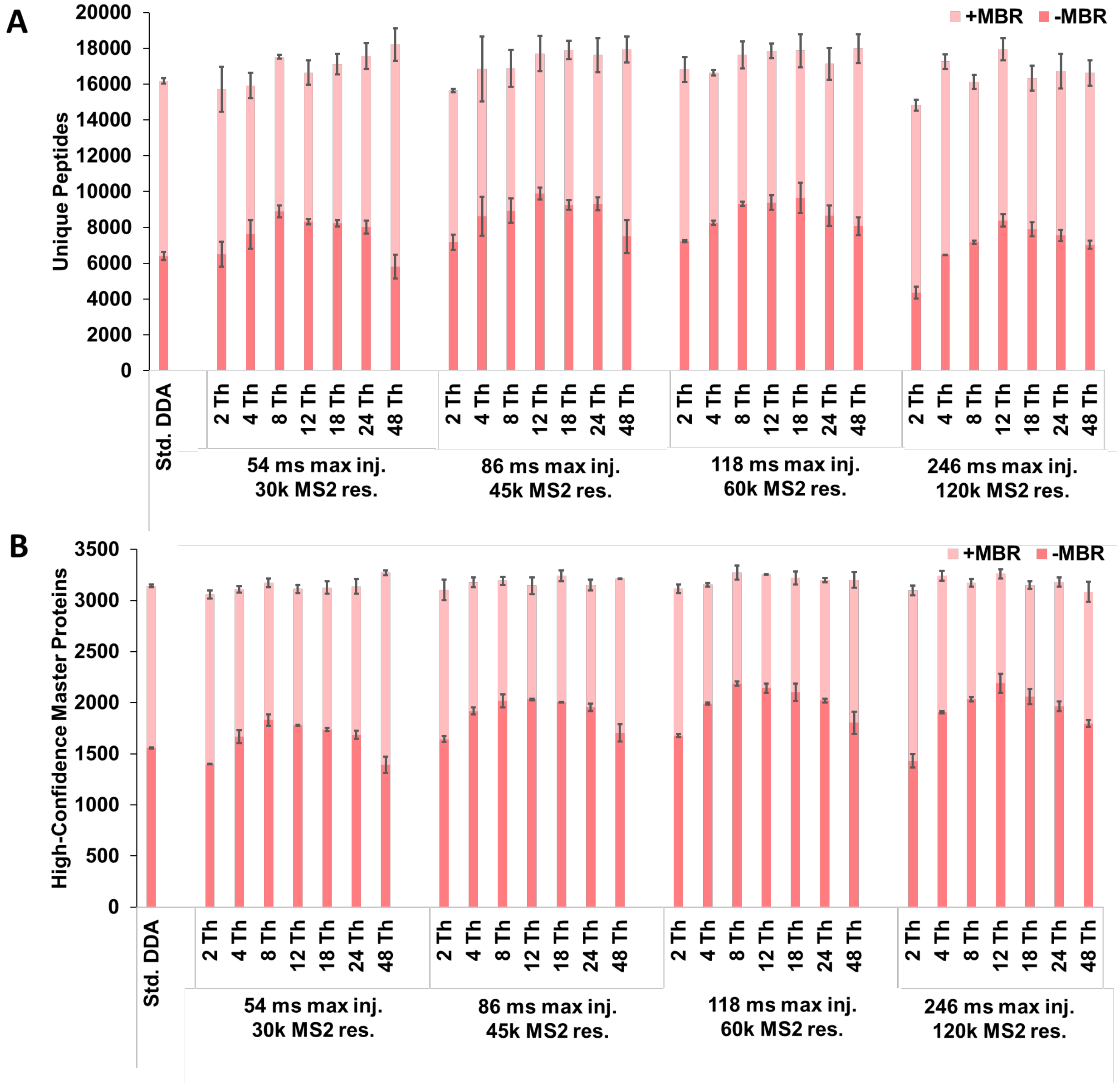


**Figure S2. Parameter optimization experiments for 20-min gradients using 0.2 ng aliquots of HeLa digest with MBR.** (A) Number of unique peptides identified as a function of MS acquisition settings. (B) Number of high-confidence master proteins (1% FDR). All identifications are based on MS2 identification and MBR. Std. DDA conditions are listed in Methods. Error bars indicate ± 1 std. dev., and n=2 for all conditions.

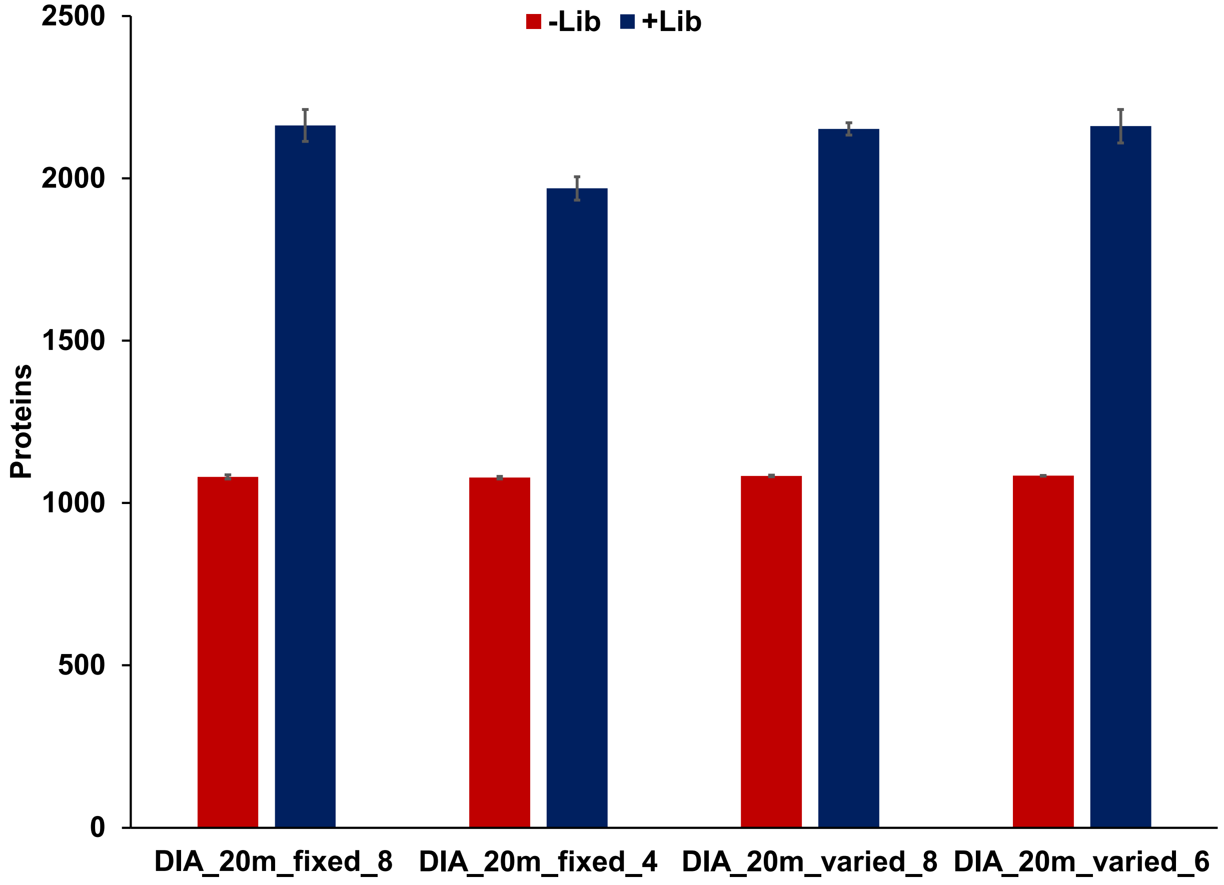


**Figure S3. Comparison between different scanning windows for 20-min gradient DIA using 0.2ng aliquots Hela digest.** Number of proteins identified as a function of scanning method with and without a spectral library. Scan windows are detailed in Supplemental Table 1.

**Supplemental Table 1. Detailed scanning windows for DIA experiments.**

| | 40-min gradient | | 20-min gradient | | | |
|---|---|---|---|---|---|---|
| | DIA_fixed_8 | DIA_varied_8 | DIA_fixed_8 | DIA_fixed_4 | DIA_varied_8 | DIA_varied_6 |
| Scan Windows | 400-450 | 400-420 | 400-450 | 400-500 | 400-420 | 400-440 |
| | 450-500 | 420-445 | 450-500 | 500-600 | 420-445 | 440-490 |
| | 500-550 | 445-475 | 500-550 | 600-700 | 445-475 | 490-550 |
| | 550-600 | 475-510 | 550-600 | 700-800 | 475-510 | 550-620 |
| | 600-650 | 510-555 | 600-650 | | 510-555 | 620-710 |
| | 650-700 | 555-605 | 650-700 | | 555-605 | 710-800 |
| | 700-750 | 605-680 | 700-750 | | 605-680 | |
| | 750-800 | 680-800 | 750-800 | | 680-800 | |
